# Corona virus fear among health workers during the early phase of pandemic response in Nepal: a web-based cross-sectional study

**DOI:** 10.1101/2020.11.04.367912

**Authors:** Pratik Khanal, Navin Devkota, Minakshi Dahal, Kiran Paudel, Shiva Raj Mishra, Devavrat Joshi

**Affiliations:** Institute of Medicine, Tribhuvan University, Kathmandu, Nepal; National Academy for Medical Sciences, Kathmandu, Nepal; Center for Research on Environment, Health and Population Activities (CREHPA), Kathmandu, Nepal; Nepal Development Society, Chitwan, Nepal

**Author notes:** **Email address of authors**, (PK); (ND); (MD); (KP); (SRM); (DJ). Corresponding author: Pratik Khanal, Institute of Medicine, 44600, Kathmandu, Nepal.

**Keywords:** COVID-19, fear, health workers, mental health, Nepal

## Abstract

**Background:** Health workers involved in COVID-19 response might be at risk of developing fear and psychological distress. This study aimed to identify factors associated with COVID-19 fear among health workers in Nepal during the early phase of pandemic.

**Methods:** A web-based cross-sectional survey was conducted in the month of April-May 2020 among 475 health workers directly involved in COVID-19 management. The Fear Scale of COVID 19 (FCV-19S) was used to measure the status of fear. Scatter plots were used to observe the relationship between fear and other psychological outcomes: anxiety, depression and insomnia. Multivariable logistic regression was done to identify factors associated with COVID fear.

**Results:** COVID-19 fear score was moderately correlated with anxiety and depression, and weakly correlated with insomnia (p<0.001). Nurses (AOR=2.29; 95% CI: 1.23-4.26), health workers experiencing stigma (AOR=1.83; 95% CI: 1.12-2.73), those working in affected district(AOR=1.76; 95% CI: 1.12-2.77) and presence of family member with chronic diseases (AOR=1.50; 95% CI: 1.01-2.25) was associated with higher odds of developing COVID-19 fear as compared to other health workers, health workers not experiencing stigma, working in non-affected district and not having family member with chronic diseases respectively.

**Conclusion:** Nurses, health workers facing stigma, those working in affect district and having family member with chronic diseases were more at risk of developing COVID-19 fear. It is thus recommended to improve work environment to reduce fear among health workers, employ stigma reduction interventions, and ensure personal and family support for those having family member with chronic diseases.

## Introduction

The psychological implications as a result of disease outbreak is often neglected by the health system[1–3] although studies have found that the proportion of mental health effects is higher than the effect of particular disease during the epidemics[4]. COVID-19 is burdening the health systems including health workforce and paralyzing economies across the world. Nepal, a South Asian country, ranking low in health security index (111 out of 195 countries[5] is not an exception from the threat of COVID-19. The country reported its first case on January 23[6] and the total infection toll has reached to 168.235 along with 920 deaths as of October 30, 2020[7]. The increasing rate of the infection is putting a strain on its already compromised health system [8]. Health care workers who are at the frontline of managing of COVID 19 are prone to developing psychological outcome as they work in a stressful situation[9]. Early evidence has shown increased work pressure, inadequate protective measures, risk of infection, and transmitting infection to family members, limited organizational support and exhaustion contributing to adverse mental outcomes including fear in health workers [3, 10–12].

Fear and stress experienced by health workers affect their work, behaviour and health outcomes [13, 14]. The understanding of fear and other psychological outcomes among health workers has not much received attention during the pandemic. There are limited published studies which have investigated the mental health impact of COVID-19 among health workers in Nepal [15, 16]. In this regard, this study aims to assess the status of COVID-19 fear among health workers involved in COVID-19 response in a low resource setting. In addition, it aims to explore the relationship of COVID-19 fear with other mental health outcomes among health workers.

## Materials and Methods

### Study design, participants and procedures

A total of 475 health workers participated in the study. A web-based cross-sectional survey was conducted among health workers directly involved in COVID-19 management in between April 26 to May 12 in 2020. Social media groups of professional organizations were identified and health workers were requested for their interest to participate in the study. Those health workers who expressed interest were personally invited to fill up the Google forms. The inclusion criteria for the study were those aged 18 years and above, currently working in Nepal, and involved in COVID-19 response. The study protocol was approved by Ethical Review Board of Nepal Health Research Council (Registration number: 2192; 315/2020).

### Measures

The fear scale of COVID 19 (FCV-19S) was used in the study for assessing the fear among health workers. It is a relatively new scale developed by Ahorsu et al in 2020 [14] and has been used in different countries including India[9], Bangladesh[17], Israel[18], Italy[19], Turkey[20] and Eastern Europe[21]. The FCV-19S has seven items and five point likert scales ranging from 1 to 5 with lower and higher value indicating strongly disagree and strongly agree respectively. The total scores ranges between 7 to 35 and the higher the score, the higher the fear of COVID-19. Similarly, the 14 item Hospital Anxiety and Depression Scale (HADS) was used for measuring anxiety (HADS-A, 7 item) and depression (HADS-D, 7 item), and 7 item Insomnia Severity Index (ISI) was used for measuring insomnia.

Socio-demographic information of the study participants was collected which included age (up to 40, >40 years), gender (male, female), ethnicity (Brahmin/Chhetri, Janajati and others), educational qualification (Intermediate and below, bachelor and masters), marital status (single, ever married), family type (nuclear and joint), profession (doctors, nurses, others), living with children (yes, no), living with older adults (yes, no), presence of chronic disease among family members (yes, no) and history of medication for mental health problems (yes, no). Similarly, work related variables included type of health facility (primary, secondary and tertiary), work experience (up to 5 and >5 years), work role in COVID-19 response (frontline, second line), adequacy of precautionary measures in work place, (not sufficient, sufficient) aware of government incentives for health workers (yes, no), perceived stigma (yes, no, do not want to answer), working in affected district (yes, no) working overtime (yes, no) and change in regular job duty during COVID-19 (yes, no). Working in affected district was defined as district with at least one case during the time of data collection.

### Data analysis

The socio-demographic and job related characteristics, and item wise response of the FCV-19S were presented in frequency and percentage. Similarly, psychometric properties of the tool were calculated and presented in S1 Table. The pattern of relationship between FCV-19S and other psychometric tools (HADS-A, HADS-D and ISI) were explored by using scatter plots and calculating correlation coefficient (Figure 1 and S2 Table). The COVID-19 fear score was categorized as presence of fear and absence of fear of COVID-19 based on the median value. Those having scored more than median (>16) were categorized as presence of fear and less than or equal to as absence of fear of COVID-19. Chi-square test was done between categorical independent and categorical dependent variable (S3 Table) and those variables significant at 10% significance level were fitted in the multivariable logistic regression model. In the regression model, the effects of gender, ethnicity, profession, education, working in affected district, family member with chronic disease, faced stigma, precautionary measures in work place, awareness about government incentive and history of medication for mental health problem was adjusted[22]. One of the independent variables, history of medication for mental health problem was also fitted into the model though it was not significant in the bivariate analysis as it was supposed to alter psychological outcomes[23]. The Variance Inflation Factor (VIF) was measured before conducting multivariable logistic regression analysis which did not detect multicollinearity (VIF value less than 1.3).

**Fig 1:**
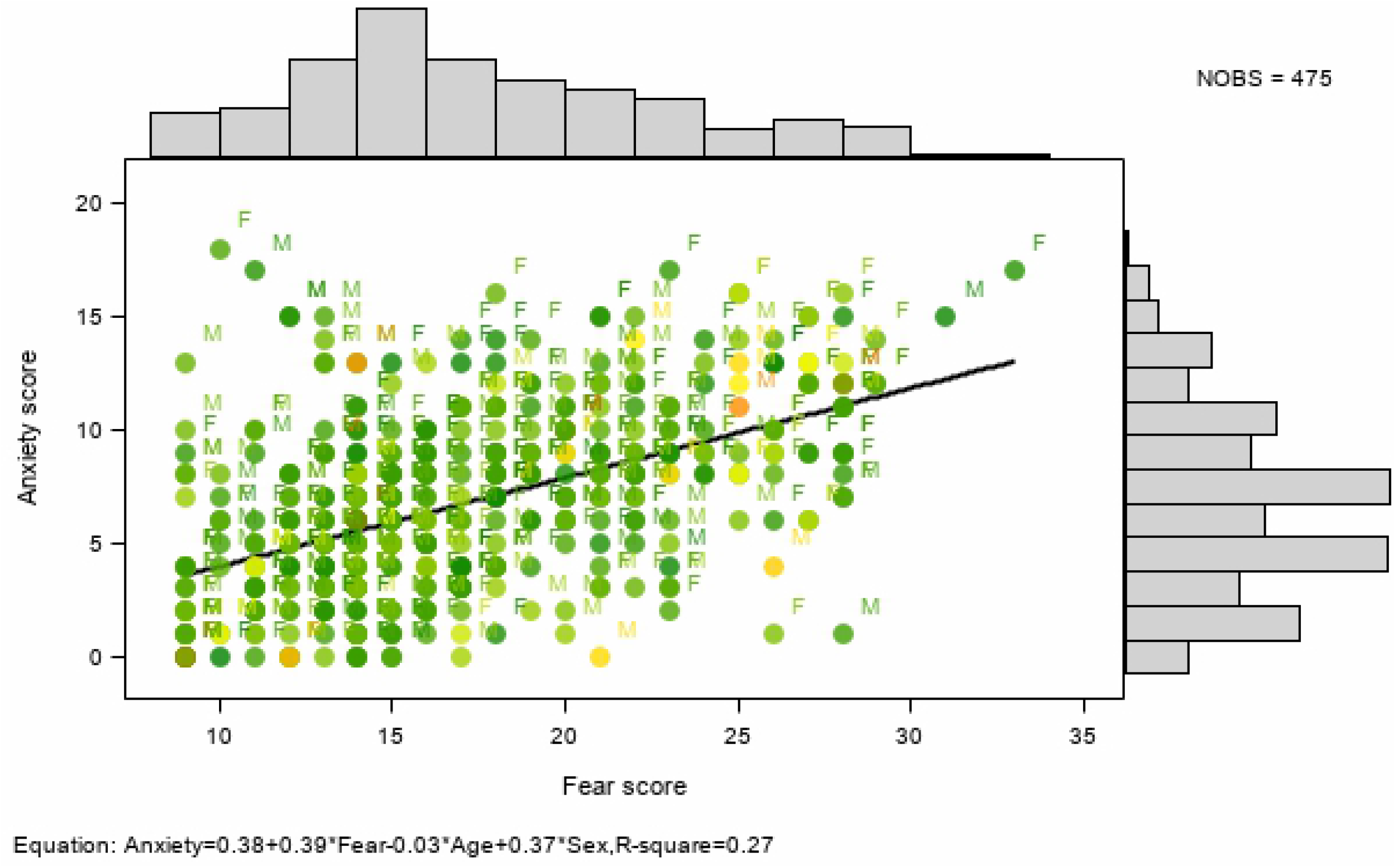
Scatter plot showing the relationship of fear, with anxiety

## Results

Tables 1 show the socio-demographic and job related characteristics of health workers. Among 475 health workers, 52.6% of them were female and 65.9% belonged to Brahmin/Chhetri ethnic group. The professional category comprised of nurses (35.2%), doctors (33.9%), paramedics (17.9%) and remaining were other health professionals. Likewise, 25.1% were living with children, 34.3% were living with elderly, 54.5% had a family member with chronic medical condition and 4.6% had a history of medication for mental health problems. Majority of the health workers in this study (82.3%) worked in either secondary or tertiary level health facility. The proportion of health workers reporting insufficient precautionary measures in the workplace, facing stigma, aware of government incentives for health workers, change in job duties during COVID-19 and working overtime was 78.9%, 53.7%, 56.8%, 70.3% and 49.1% respectively.

**Table 1:**
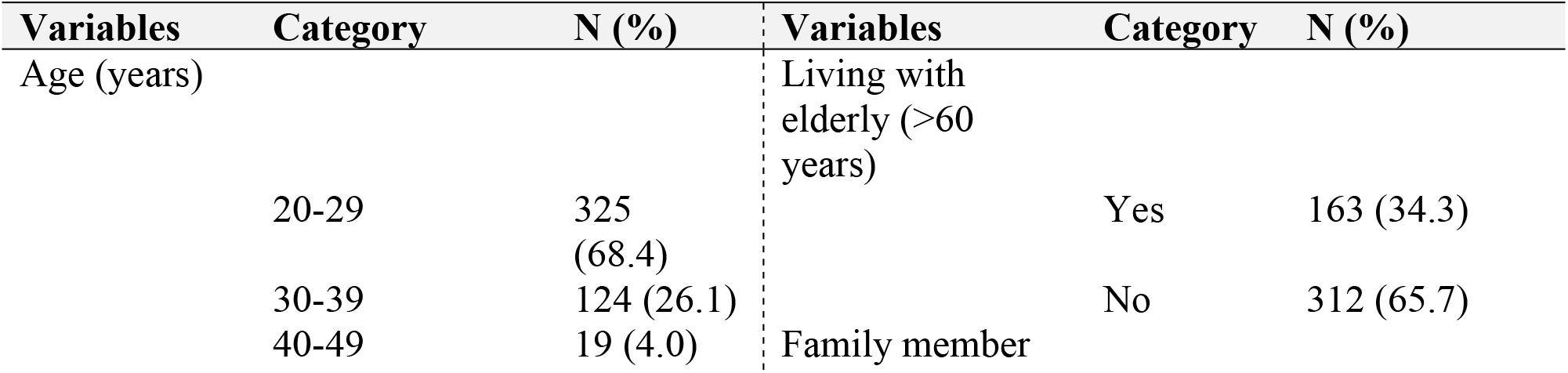
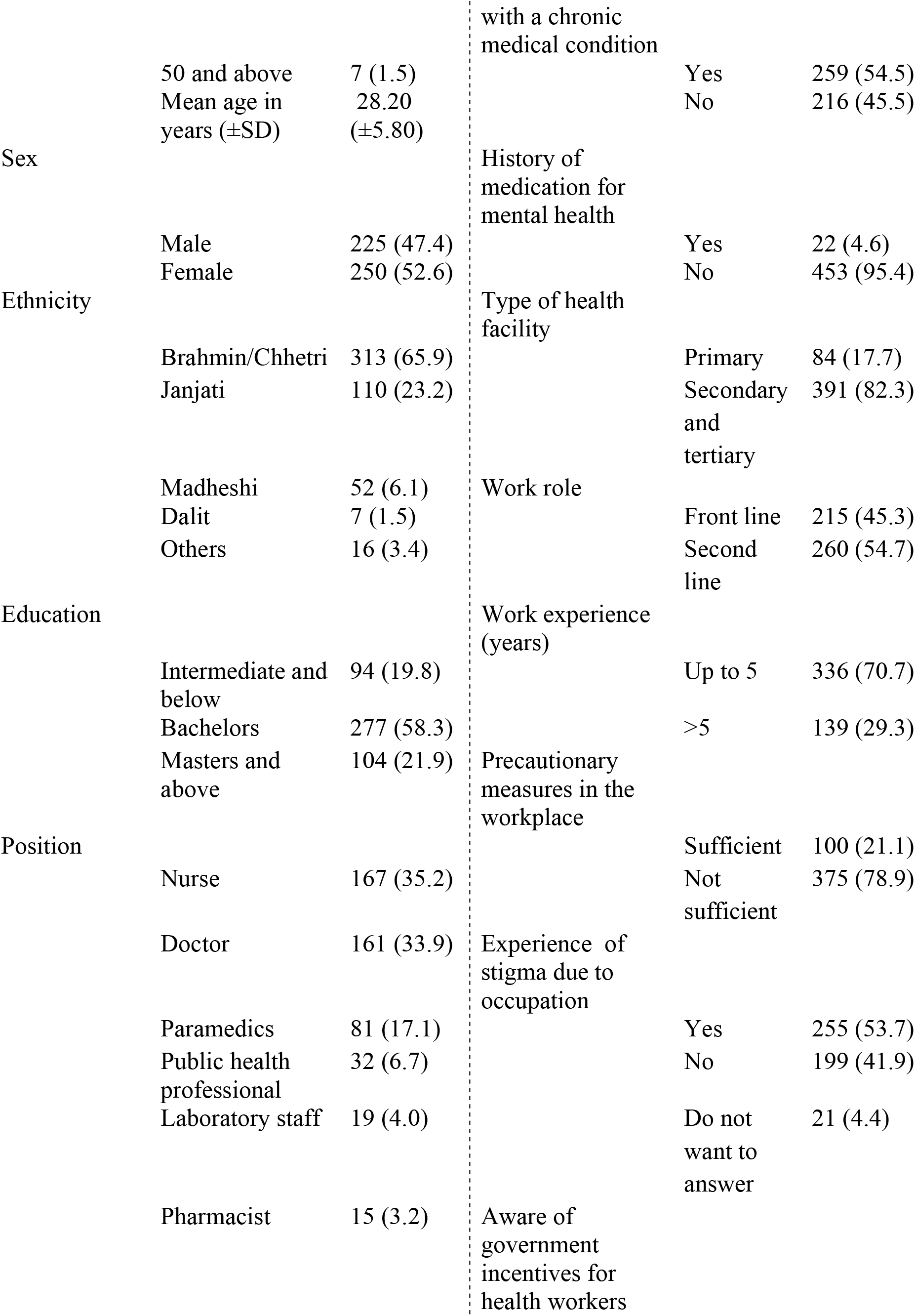
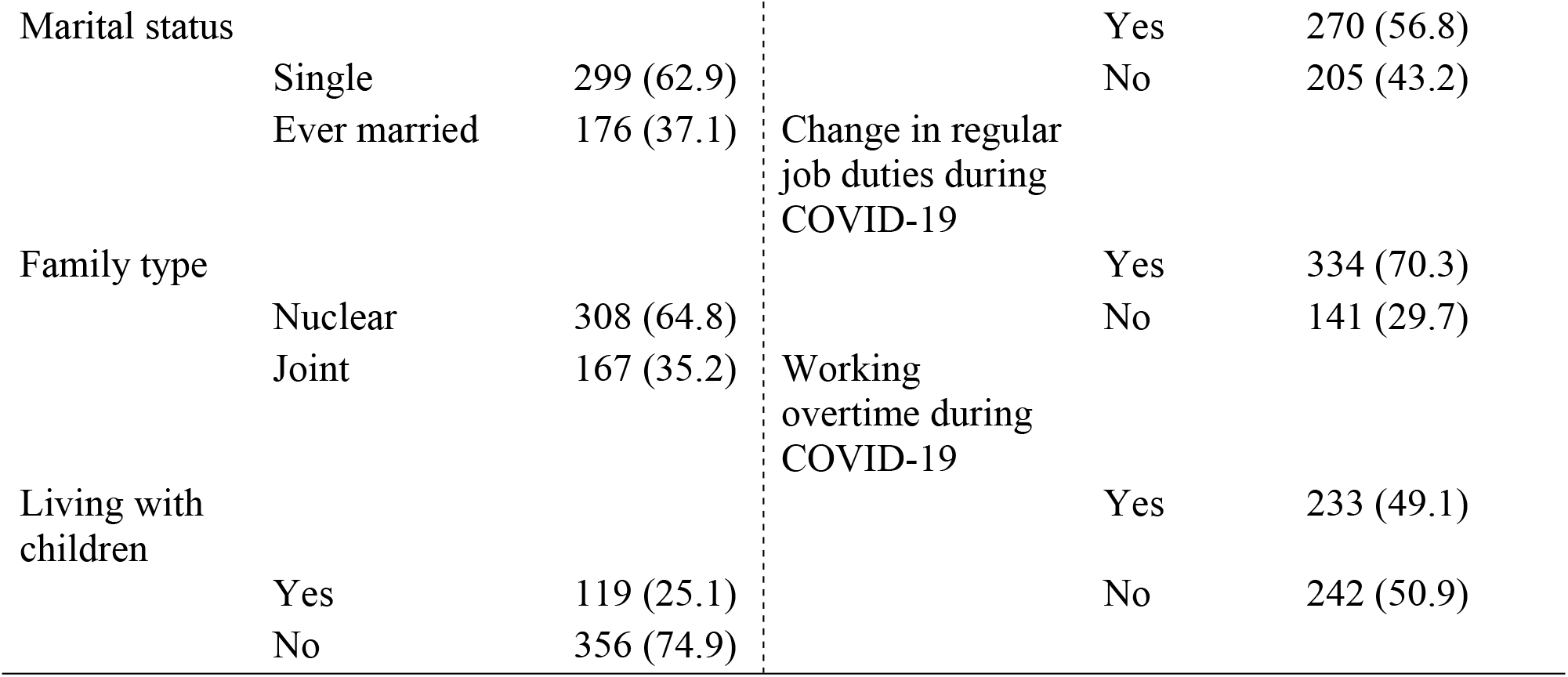
Socio-demographic and job related characteristics of health workers

The Table 2 shows the item-wise distribution of responses of FCV-19S respectively. The proportion of health workers who either strongly agree or agree to the individual items of FCV-19S was highest (32.5%) for ‘When watching news and stories about corona on social media, I become nervous and anxious’ and lowest (7.3%) for ‘I cannot sleep because I am worrying about getting Corona’.

**Table 2:** Item-wise distribution of responses

| Scale | Items | Strongly disagree | Disagree | Neither agree nor disagree | Agree | Strongly agree |
| --- | --- | --- | --- | --- | --- | --- |
|  |  | N (%) | N (%) | N (%) | N (%) | N (%) |
| FCV-19 S1 | I am most afraid of corona virus disease-19 | 65 (13.7) | 150 (31.6) | 132 (27.8) | 103 (21.7) | 25 (5.3) |
| FCV-19 S2 | It makes me uncomfortable to think about corona | 80 (16.8) | 177 (37.3) | 84 (17.7) | 122(25.7) | 12 (2.5) |
| FCV-19 S2 | My hands become clammy when I think about corona | 159 (33.5) | 196 (41.3) | 74 (15.6) | 40 (8.4) | 6 (1.3) |
| FCV-19 S4 | I am afraid of losing my life because of corona | - | 304 (64.0) | 77 (16.2) | 80 (16.8) | 14 (2.9) |
| FCV-19 S5 | When watching news and stories about corona on social media, I become nervous and anxious | 87 (18.3) | 150 (31.6) | 84 (17.7) | 129 (27.2) | 25 (5.3) |
| FCV-19<br>S6 | I cannot sleep because I am<br>worrying about getting<br>Corona | - | 372 (78.3) | 68 (14.3) | 31(6.5) | 4 (0.8) |
| FCV-19<br>S7 | My heart races or palpitates<br>when I think about getting<br>corona | 147(30.9) | 192 (40.4) | 76 (16.0) | 48 (10.1) | 12 (2.5) |

### Correlation of FCV-19 S with HADS-A, HADS-D and ISI

The correlation analysis showed that FCV-19S was moderately correlated with HADS-A (r= 0.513, p<0.001) and HADS-D (r= 0.425, p<0.001) while weakly correlated with ISI (*r*= 0.367, p<0.001). The seven items of the FCV-19S were either weakly or moderately correlated with HADS-A, HADS-D and ISI (p<0.001) (S2 Table). The scatter plot showing the relationship between anxiety and fear, depression and fear, and insomnia and fear adjusted for age and sex is shown in Figure 1, Figure 2 and Figure 3 respectively.

**Fig 2:**
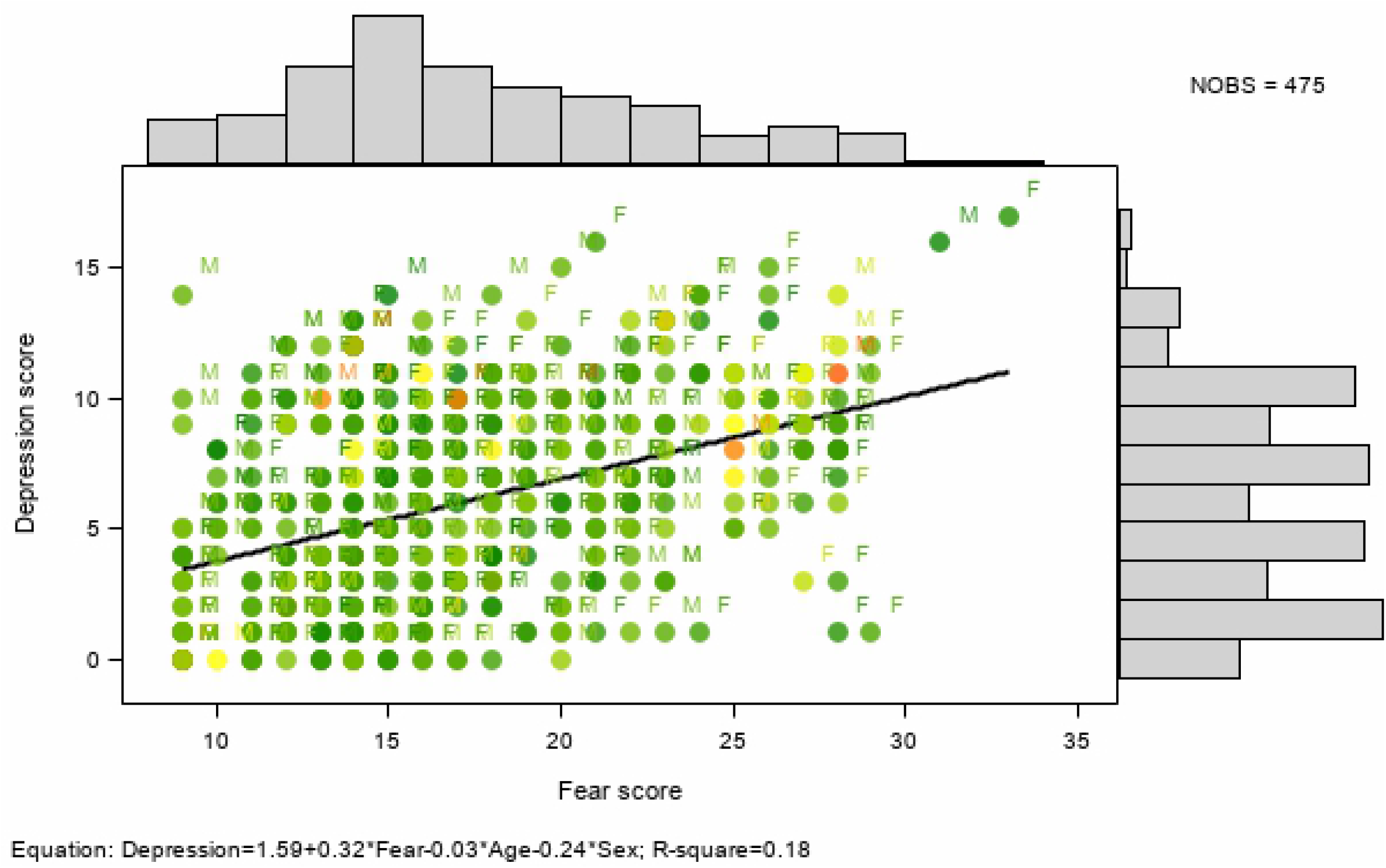
Scatter plot showing the relationship of fear, with depression

**Fig 3:**
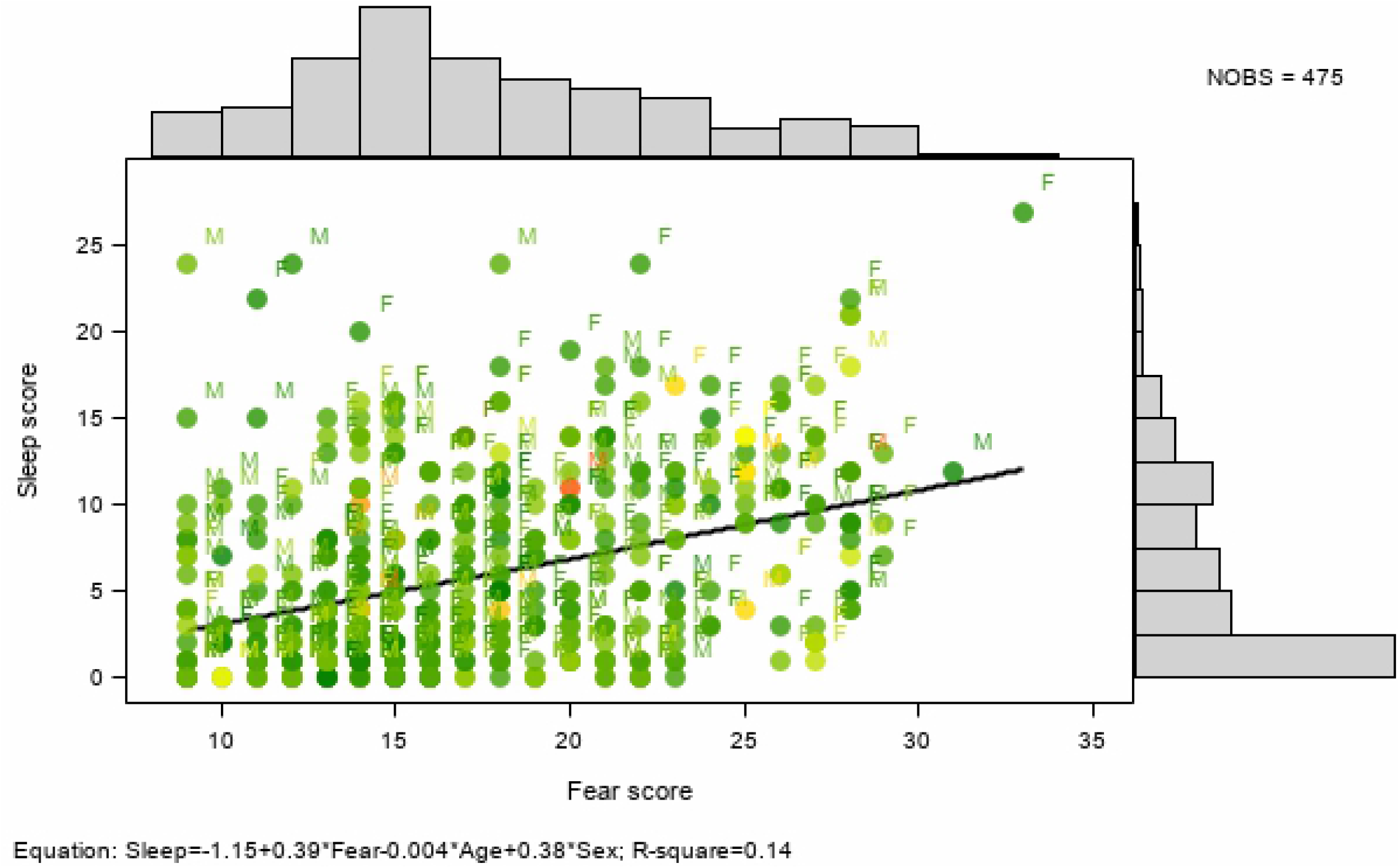
Scatter plot showing the relationship of fear, with insomnia. The colour response is based in age (years) using a colour-gradient: from green, yellow to red for lowest to the highest age. The equation in the footnote shows the relationship of fear with anxiety, depression and insomnia adjusted for age and sex

### Predictors of COVID-19 fear among health workers

The bivariate analysis between socio-demographic and job related characteristics is presented in S3 Table. The proportion of COVID-19 fear among health workers in this study was 46.1 % (219/475). In the adjusted analysis, profession, stigma experience, working in affected district and having family member with chronic disease was significantly associated with COVID fear. As compared to other health workers, nurses (AOR=2.29; 95% CI: 1.23-4.26) had significantly higher odds of having COVID fear. Similarly, health workers working in affected district (AOR=1.76; 95% CI: 1.12-2.77), those having family member with chronic disease (AOR=1.50; 95% CI: 1.01-2.25), and those who faced stigma (AOR=1.83; 95% CI: 1.12-2.73) had significantly higher odds of having COVID fear as compared to those not working in affected district, not having a family member with chronic disease, and those not facing stigma respectively. Gender, ethnicity, education, precautionary measures, awareness about government incentive and history of medication for mental health problems was however not statistically significant with COVID fear (Table 3).

**Table 3:**
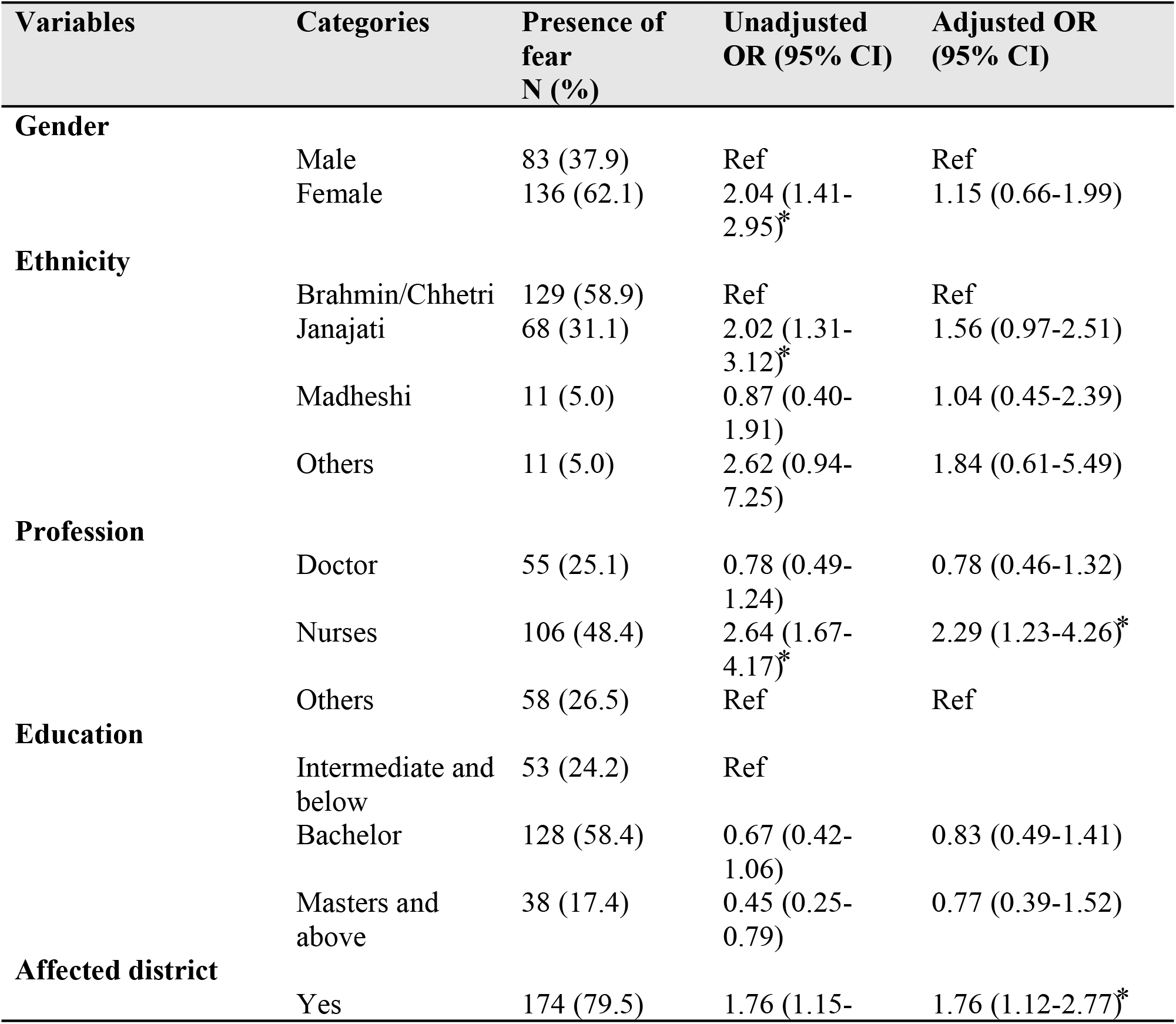
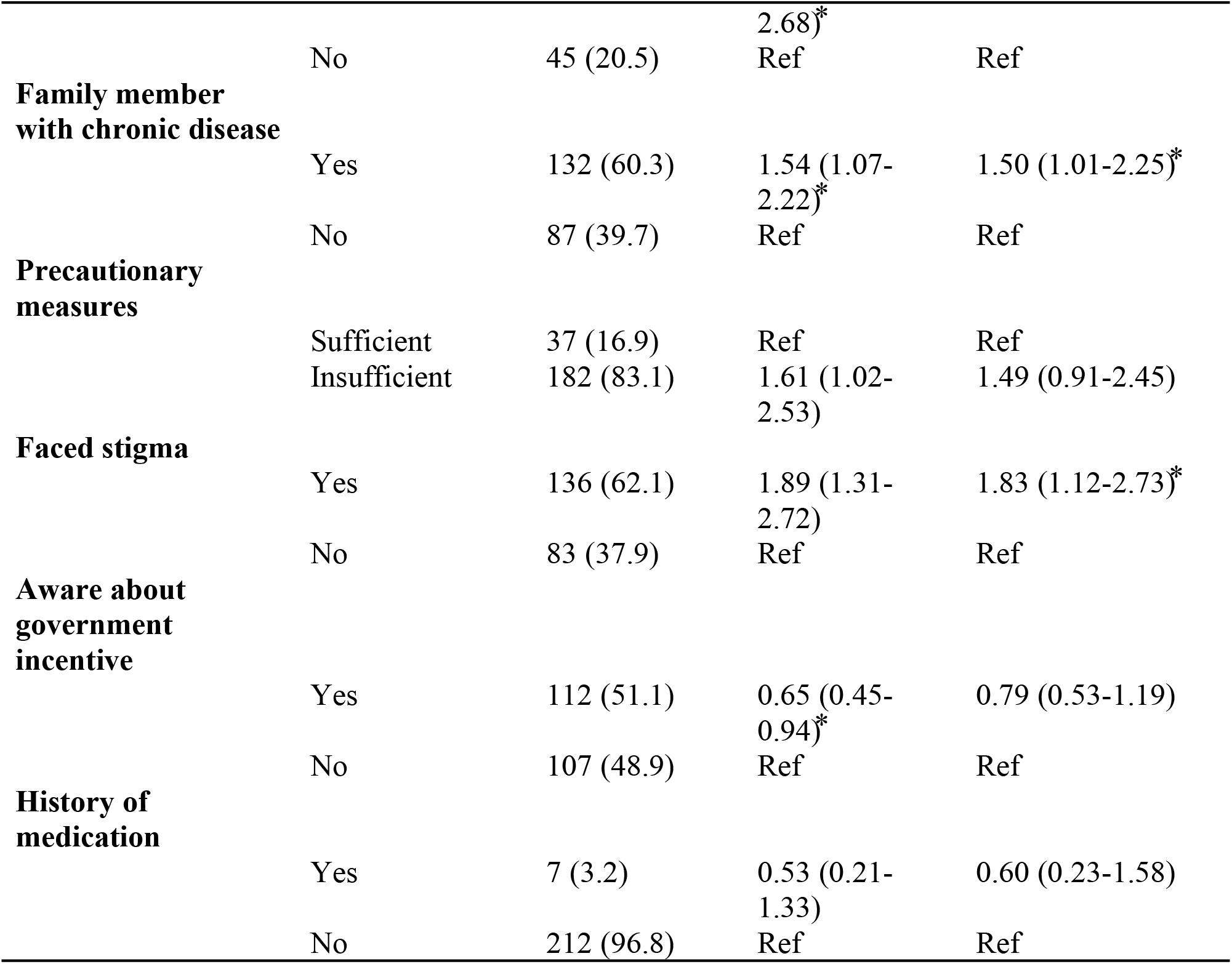
Factors associated with COVID related fear among health workers (n=475)

## Discussion

This study documents the factors associated with the presence of fear related to COVID-19 among health workers in Nepal in the early phase of the pandemic. The study identified profession, working in the affected region, presence of family member with chronic disease and stigma faced by health workers as significant factors contributing to the presence of fear among health workers. In this study, nurses were significantly more likely to have COVID fear than other health workers. This might be because of their role in providing patient care more closely, frequently and for longer hours compared to other health workers. The chance of being infected and transmitting infection to others, dealing with the disease that is highly infective and the uniqueness of the cases might have led to increased fear among nurses. Similar findings were noted in studies conducted in other countries that have reported COVID-19 cases and countries that have handled epidemics like SARS in the past [24–27]. Effective strategies to reduce fear with focus on nurses are thus required to avert COVID fear and psychological distress.

In our study, more than half of the health workers experienced stigma during COVID-19. Stigma faced by health workers was also significantly associated with the higher odds of presence of fear of COVID-19. Already vulnerable due to exposure to possible infections, emotional exhaustion due to increasing workload, deployment to newer settings like fever clinics and lack of adequate PPEs, health workers are more likely to face stigma either internalized or from public which will impair their performance in COVID-19 response[28]. Stigma reduction strategies should thus be employed for educating the public which need to include proper messaging through mass media and community engagement activities[29, 30]. Equally important is to identify the underlying causes of stigma experienced by health works during the epidemic.

Working in the affected district was significantly associated with the presence of fear among health workers. This is obvious as health workers working in the affected districts need to directly deal with COVID-19 patients or those at risk of infection. Health workers in Hubei province of China[24] during COVID pandemic and health workers directly involved in the care of patients in Canada[31] during SARS epidemic also faced more psychological distress as compared to those not involved in the direct care of COVID patients or less affected areas. As fear among health workers reflects their psychological wellbeing, health workers working in risk districts should be supported emotionally and due attention is required on their workload, safety needs and other personal and family concerns.

In this study, presence of family member with chronic disease had higher odds of presence of COVID-19 fear. The fear of transmission to family members and the vulnerability posed by chronic disease conditions might have resulted in higher degree of fear among health workers. This finding is similar to the study from China[32] where health workers were concerned with the infection of their family members. Personal and family support is thus required for health workers having family member with chronic diseases.

Our study findings showed COVID fear was moderately correlated with anxiety and depression suggesting detrimental effect of COVID fear to psychological well-being. Perhaps, symptoms of anxiety and depression were a consequence of working in a high fear environment for an extended period. It is thus necessary to develop an enabling work environment where health workers feel protected and are motivated to confront COVID-19 and other similar epidemics. Health facility managers need to monitor the psychological well-being of their staffs and ensure proper psychological intervention measures are adopted timely and precisely. In this study, only one out of five health workers mentioned protective measures in their workplace as sufficient. Similarly, just over a half of health workers were aware of the government incentives entitled to them during COVID-19. This clearly reflects the need to improve organizational and policy aspects for boosting the work morale of health workers to reduce fear and psychological distress among health workers involved in COVID-19 response.

Majority of the socio-demographic and job related characteristics including work role, precautionary measures in the work place, working overtime and awareness regarding incentives were not significantly associated with the fear of COVID-19. Further follow-up studies might be required among health workers to understand the effect of socio-demographic and job related characteristics in psychological outcome such as fear.

Our study has some limitations to be noted. This study was conducted during the early phase of the pandemic in Nepal when less than 300 COVID-19 cases were reported. The status of fear might have altered thereafter as COVID-19 cases continue to increase in Nepal. Similarly, participation in this study required internet access and the survey was administered in English language. This might have left out health workers who did not have internet access and had difficulty in comprehending English language. Similarly, the results might have been affected by subjective response. The feeling of uncertainty about the scale and duration of the epidemic, no known medication or vaccine, widespread media coverage and news about surge of cases and deaths in various affluent countries with sophisticated health system and lack of adequate testing facilities might also have accentuated the perceived level of fear among healthcare workers. Despite limitations, this study employs FCV-19S to measure the status of fear among health workers and identify those at risk of developing fear. The evidence generated can be useful to those at decision making level and health facility managers for designing appropriate interventions to enhance psychological well-being among health workers in this and similar epidemics in the future.

## Conclusion

This study showed a considerate proportion of COVID fear among health workers during the early phase of pandemic in Nepal. Nurses, health workers working in affected district, those facing stigma and having family member with chronic diseases were significantly more likely to have COVID fear than other health workers, health workers working in non-affected district, those with no stigma experience and those not having a family member with chronic disease. Based on the study findings, it is recommended to focus on strategies to improve work environment to reduce fear among health workers, conduct stigma reduction activities, and ensure personal and family support for health workers having family member with chronic diseases.

## List of abbreviations

AOR: adjusted odds ratio
CI: confidence interval
COVID-19: Corona virus 2019
FCV-19S: Fear of COVID-19 Scale
HADS: Hospital Anxiety Depression Scale
ISI: Insomnia severity index
PPE: personal protective equipment
SARS: Severe Acute Respiratory Syndrome

## Acknowledgements

The authors would like to acknowledge all the health workers involved in the study and the Policy, Planning and Monitoring Division of Ministry of Health and Population for providing us a letter of support for the study.

## Author’s contribution

Conceptualization: Pratik Khanal, Navin Devkota, Kiran Paudel

Data curation: Navin Devkota, Kiran Paudel

Formal analysis: Pratik Khanal, Minakshi Dahal, Shiva Raj Mishra

Methodology: Pratik Khanal, Navin Devkota, Kiran Paudel, Devavrat Joshi

Supervision: Devavrat Joshi

Writing-original draft: Pratik Khanal, Navin Devkota, Minakshi Dahal, Kiran Paudel

Writing-review and editing: Shiva Raj Mishra, Devavrat Joshi

## Supporting Information

S1: Descriptive analysis of the items of the English version FCV-19S (Doc)

S2: Correlation of FCV-19 S with HADS-A, HADS-D and ISI (Doc)

S3: Fear of COVID-19 and its associated factors (Doc)

